## Supplemental file for "Dissociable memory modulation mechanisms facilitate fear amnesia at different timescales"

**Supplemental Table S1**. Participant exclusion criteria.

|  | **Total subject #** | **Non-responder** | **Non-learner** in **Acquisition** | **Non-learner** in **Extinction** | **Final subject #** |
| --- | --- | --- | --- | --- | --- |
| Exclusion criterion |  | Mean CS SCR < 0.02 u*S* | 1. Mean CS+ SCR < CS- SCR in the latter half trials AND 2. diff SCR (CS+, CS-) in the 2^nd^ half trials **<** diff SCR (CS+, CS-) the 1^st^ half trials | 1. Mean CS+ SCR > CS- SCR in the latter half and last trials) AND 2. diff SCR (CS+, CS-) 2^nd^ half trials **>** diff SCR (CS+, CS-) the 1^st^ half trials |  |
| Study 1 (reminder group) | 36 | 0 | 3 | 3 | 30 |
| Study 1 (no-reminder group) | 39 | 8 | 1 | 3 | 27 |
| Study 2 (30min group) | 48 | 19 | 1 | 1 | 27 |
| Study 2 (6hr group) | 31 | 5 | 0 | 0 | 26 |
| Study 2 (24hr group) | 40 | 13 | 1 | 0 | 26 |
| Study 3(R-PFC) | 22 | 3 | 0 | 0 | 19 |
| Study 3 (R-VER) | 22 | 4 | 0 | 0 | 18 |
| Study 3 (NR-PFC) | 22 | 4 | 0 | 0 | 18 |
| Study 3(NR-VER) | 24 | 4 | 0 | 0 | 20 |

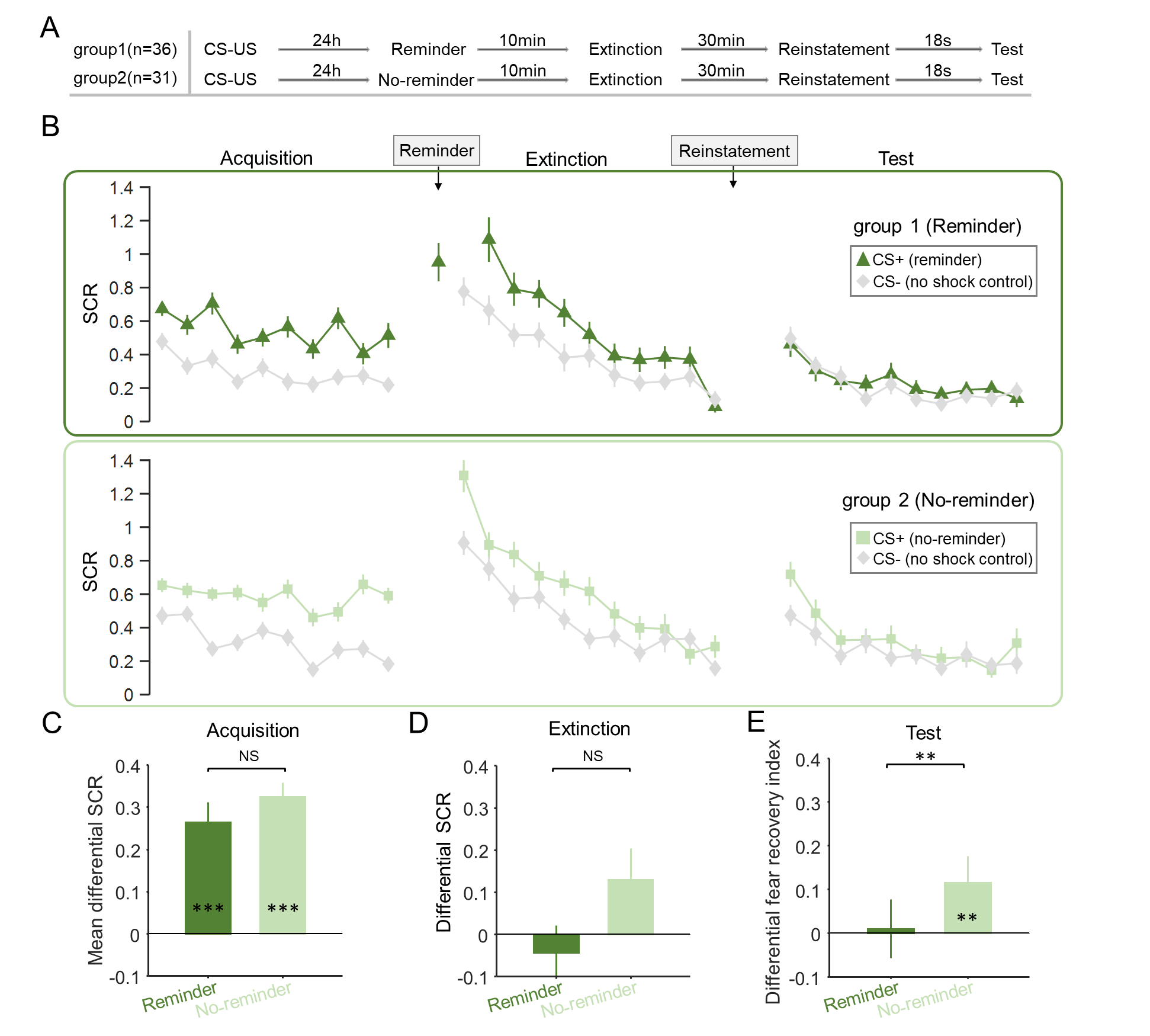

**Supplemental Figure 1. SCR responses to conditioned stimuli in two groups of study 1 with all responders (learners + non-learners).** (**A**) Experimental design and timeline. (**B**) Mean SCRs of fear conditioned stimuli (CS+) and the control stimulus (CS-) across fear acquisition, extinction and test phases for two groups (reminder and no-reminder). (**C**) Mean differential SCRs (CS+ minus CS-) in the acquisition phase (latter half trials). (**D**) Mean differential SCRs (CS+ minus CS-) in the extinction phase (last trial). (**E**) Differential fear recovery index (FRI) for the reminder and no-reminder groups (CS+ FRI minus CS- FRI). ****P* < 0.001, ***P* < 0.01. NS: Non-significant. Mann-Whitney U test, error bars represent standard errors.

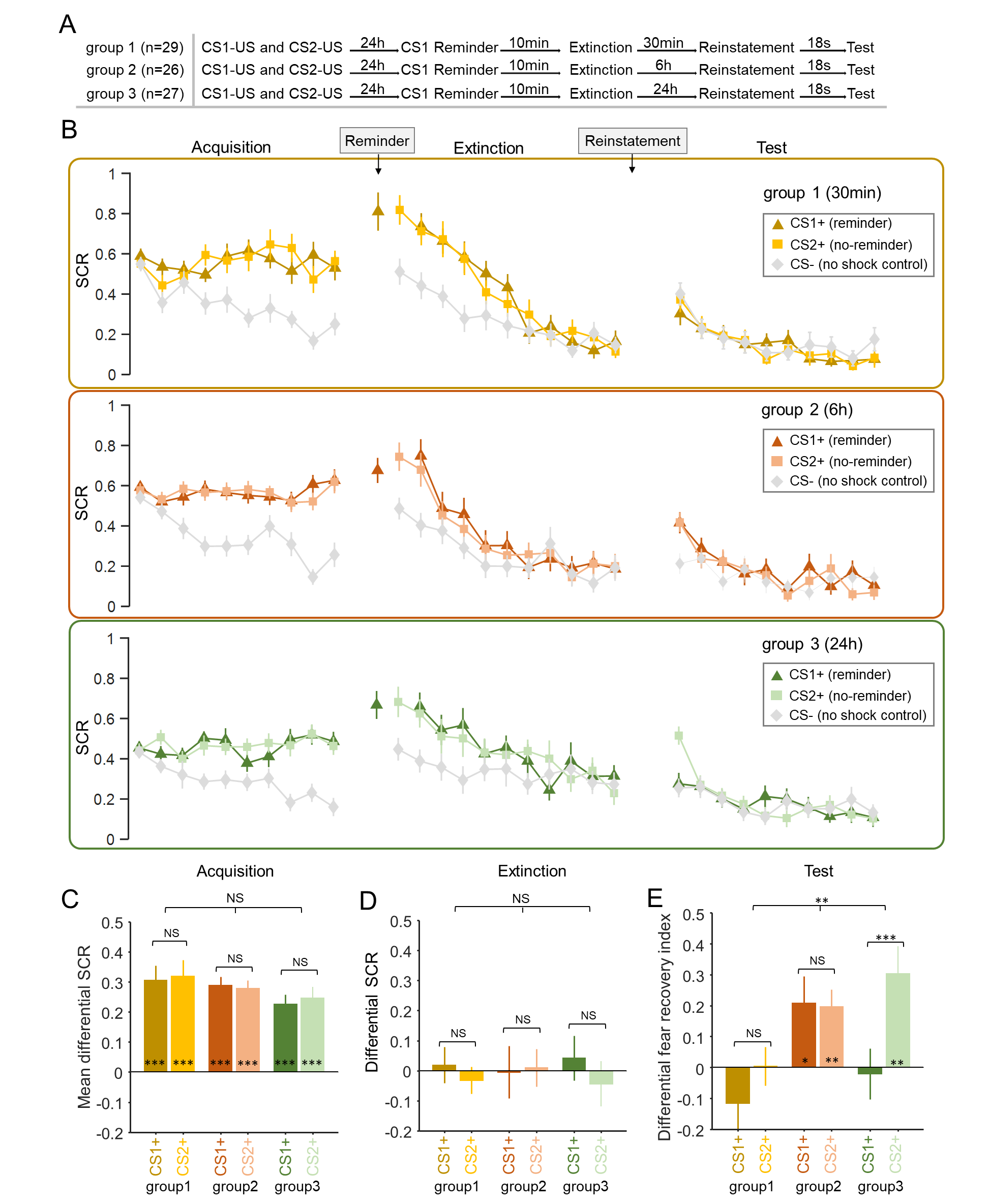

**Supplemental Figure 2. SCR responses to conditioned stimuli in three groups of study 2 with all responders (learners + non-learners).** **(A)** Experimental design and timeline. **(B)** SCRs of fear conditioned stimuli CS1+ (reminder) and CS2+ (No-reminder), and the control stimulus (CS-) across the fear acquisition, extinction and test phases for each group (30min, 6h and 24h tests). **(C)** Mean differential SCRs (CS+ minus CS-) in the acquisition phase (latter half trials). **(D)** Differential SCRs (CS+ minus CS-) in the extinction phase (last trial). **(E)** Differential fear recovery index (FRI) for the reminder and no-reminder groups (CS+ FRI minus CS- FRI). ****P* < 0.001; **P* < 0.05. NS: Non-significant. Error bars represent standard errors.

For the learners in study 1, since the SCRs of CS+ and CS- were not significantly different in the last extinction trial, we then focused on the test phase and performed a mixed two-way ANOVA on the first test trial SCR with the group (reminder vs. no-reminder) and CS (CS+ vs. CS-) factors and it showed a significant CS × group interaction (*F*_1,55_ = 8.737, *P* = 0.005, *η*^2^ = 0.137). Post hoc *t*-tests showed that SCRs invoked by the CS+ were significantly larger than that of the CS- in the test phase (first trial, *t*_26_ = 4.032, *P* < 0.001) for the no-reminder group, whereas SCRs were not significantly different between CS+ and CS- in the reminder group (*t*_29_ = -0.394, *P* = 0.696). Importantly, the differential SCRs (difference between CS+ and CS-) for the CS+ in the reminder and no-reminder groups are statistically different (*t*_55_ = -2.956, *P* = 0.005).

For all the learners in study 2, since the SCRs of the CS+ and CS- were not statistically different in the last extinction trial, we also performed a mixed two-way ANOVA with the within-subject factor CS (CS-, CS1+ and CS2+) and the between-subject factor group (30min, 6h and 24h) on the first test trial SCR and found a significant interaction effect on all learners (*F*_4,152_ = 11.611, *P* < 0.001, *η*^2^ = 0.234). We further examined the effects in each group. In the 30min group, post-hoc *t*-tests showed that the retrieval-extinction training diminished fear responses to both the reminded conditioned stimulus (CS1+ vs. CS- SCR, *t*_26_ = -1.790, *P* = 0.085) and the non-reminded CS+ (CS2+ vs. CS- SCR, *t*_26_ = -0.495, *P* = 0.625), suggesting a cue-independent short-term amnesia effect. In addition, there was no significant difference between the SCRs of CS1+ and CS2+ (*t*_26_ = -1.243, *P* = 0.225).

In contrast to the short-term effect, in the 6h group, post-hoc *t*-tests showed that fear memory recovered for the reminded CS+ (CS1+ vs. CS- SCR, *t*_25_ = 5.496, *P* < 0.001), as well as the non-reminded CS+ (CS2+ vs. CS- SCR, *t*_25_ = 5.670, *P* < 0.001) in the medium-term test of the retrieval-extinction procedure effect, suggesting the failure to suppress the return fear memory in the 6h group. There was also no significant difference between SCRs between CS1+ and CS2+ (*t*_25_ = -0.125, *P* = 0.901).

Finally, in the 24h group, the retrieval-extinction procedure only diminished fear responses to the reminded CS+ (CS1+ vs. CS- SCR, *t*_25_ = 0.402, *P* = 0.691), whereas the fear response to the non-reminded CS+ remained significant (CS2+ vs. CS- SCR, *t*_25_ = 4.827, *P* = 0.003). Post-hoc t-test showed that the SCRs of CS1+ and CS2+ were significantly different (*t*_25_ = -4.577, *P* < 0.001).
